# Seronegative MSM at high-risk of HIV-1 acquisition show Immune quiescent profile with normal immune response against common antigens

**DOI:** 10.1101/2021.03.11.434877

**Authors:** Ana C. Ossa-Giraldo, Yurany Blanquiceth, Lizdany Flórez-Álvarez, Katherin Contreras-Ramírez, Mauricio Rojas, Juan C. Hernandez, Wildeman Zapata

## Abstract

HIV infection still represents a major public health problem worldwide, and a vaccine remains elusive. The study of HIV-exposed seronegative individuals (HESN) brings important information about the natural resistance to HIV, allows a better understanding of the infection and opens doors for new preventive and therapeutic strategies. Among HESN groups there are some men who have sex with men (MSM) with high-risk sexual behaviors, who represent an adequate cohort for the study of HESN because of their major exposure to HIV in the absence of infection. This study aimed to compare the immunological profile of Colombian seronegative MSM with different risk sexual behaviors. Sixty MSM at high-risk (n=16) and low-risk (n=44) of HIV-1 acquisition were included. No sex worker nor homozygous delta 32 mutation subjects were included. All the participants were negative for anti-HIV-1/2 antibodies and HIV-1 proviral DNA. The high-risk MSM presented a higher frequency of sexual partners in the last 3 months previous to the study’s enrollment (Median 30 vs. 2), lifetime sexual partners (Median 1708 vs. 26), and unprotected anal intercourse (Median 12.5 vs. 2) than low-risk MSM. This group also showed a quiescent profile of T cells and NK cells, with a significantly lower percentage of CD4+CD38+, CD4+HLADR-CD38+, CD4+Ki67+ T cells, NKG2D+ NK cells (CD3-CD16+CD56+), a significantly higher percentage of CD4+HLADR-CD38- and a tendency to show a higher percentage of CD8+HLADR+CD38-T cells, than the low-risk group. Likewise, they showed higher mRNA levels of Serpin A1 from PBMCs. The results suggest that this cohort of MSM could be HESN individuals and their resistance would be explained by a quiescent profile of T cells and NK cells, and increased expression of Serpin A1. It is necessary to continue the study of MSM at high-risk of exposure to HIV-1 to better understand the natural resistance to HIV.

## Introduction

HIV-1 infection remains to be a big health issue and priority worldwide, despite the availability of highly effective antiretroviral therapy (1). Several efforts to find a vaccine have been done but it continues elusive (2). Some people -HIV-exposed seronegative individuals (HESN)-exhibit a natural resistance to HIV-1, who persist without an established infection besides their exposure to the virus (3). The comprehension of HESN’s biological phenomena is an advantage to better understand the HIV-1 infection and brings new options for therapeutic and preventive strategies design (3,4). Since the first reports of natural resistance to HIV-1 (5,6), the mechanisms that can explain this phenomenon in HESN individuals are a major concern for HIV researchers, and still current now.

The main factors associated with HIV natural resistance are the homozygous deletion of 32bp in CCR5 co-receptor or Δ32 mutation, present in near to 1% of the global population (7), and recently described, the heterozygous single nucleotide deletion in the stop codon of the nuclear import factor Transportin 3 gene (*TNPO3*) (8). However, not all HESN subjects have these protective genotype and other immunological and genetic factors have been associated with HIV-1 natural resistance in HESN cohorts, such as the overexpression of antimicrobial peptides in the mucosa, increased activity of NK cells, a combination of certain HLA-KIR genotypes, and HIV-1-specific CTL (cytotoxic T cells) responses (9–13).

In the context of HIV-1 exposure the presence of activated cells and inflammation at the site of HIV entry increase the risk of infection, as reported in people with clinical or subclinical STIs and mucosal inflammatory process (14–17). The immune quiescence, characterized by a low activation profile of target cells, has been proposed as a protection mechanism against HIV infection and it has been found in several HESN cohorts, including men who have sex with men (MSM) (18–22). In previous studies in the Pumwani cohort, a well-studied HESN cohort of African female sex workers, reduced susceptibility to HIV-1 infection was related to a lower frequency of activated CD4+CD69+ T cells and an elevated number of Treg cells (23). Low expression of genes implicated in T cells receptor signaling and HIV host-dependent factors also have been described together with lower levels of secreted cytokines at basal state, but no after stimulation, showing an important difference between quiescence and immunosuppression (24,25). Contrariwise, other studies have shown that the immune activation state is related to protection. Biasin *et al*. reported in 2000 an increased expression of proinflammatory cytokines and chemokine receptors in cervical biopsies of HESN women (26). Increased memory and activated T cells have been also described (27,28). These results show that the role of cell activation in HIV-1 protection is not clear.

Since the first reports of resistance in 1989 (6) to the present, MSM have represented an adequate cohort for the study of HESN individuals. The high prevalence of HIV in this population (1) plus the higher probability to acquire the infection through anal sex (29), put MSM who practice risky sexual behaviors, such as anal sex without a condom and having multiple partners, at extreme risk of HIV exposure. Therefore, the study of seronegative MSM with high-risk behaviors and the possible finding of HESN individuals in this population represents an important opportunity to better understand HIV natural resistance.

The HIV prevalence in Colombian MSM is ∼43 times higher than in the general population [17% vs. 0.4%] (1), and pre-exposure prophylaxis (PrEP) has not yet been approved in the country. Therefore, Colombian seronegative MSM that practice high-risk sexual behaviors are a very interesting population to address the mechanisms underlying HIV natural resistance. However, engaging this population in Latin America is not easy because most MSM do not want to share private information or even being identified due to the historic oppression they have suffered. This study aimed to compare the immunological profile of Colombian HIV seronegative MSM.

## Material and methods

### Study population

Sixty subjects were included from a cohort of MSM from Medellin-Colombia who signed informed consent and accepted to participate in the study. The recruitment of participants was carried out by the combination of several methods to sample hard-to-reach populations and sociodemographic data and sexual behaviors were defined by structural surveys and in-depth interviews, as we previously reported (30). The subjects were classified into two groups according to the frequency of sexual partners in the last three months before the inclusion of the study. The high-risk group (n=16) was defined as MSM with more than 14 sexual partners, and the low-risk group (n=44) were MSM with four or fewer sexual partners in the last three months, respectively. All individuals met the following inclusion/exclusion criteria: no one was a sex worker nor taking PrEP; all of them were negative for anti-HIV-1 antibodies, HIV-1 proviral DNA, and delta 32 mutation in the *CCR5* gene in a homozygous state. The study was performed according to the Helsinki declaration and was approved by the Ethics Committee of Universidad de Antioquia’s School of Medicine (Act No.007, May 22th, 2014).

### Biological Samples

Peripheral blood and anal mucosa samples were obtained from each subject; the whole blood was used to obtain plasma and serum, to extract DNA, as well as for PBMCs isolation. The anal mucosa sample was used for cytology analysis and mRNA extraction and collected using an optimized protocol: the cytobrush was inserted 5 cm beyond the anal verge, close to the anal wall, rotated slowly while were withdrawn to capture cells. Then, the sample was spread onto slides for cytology analysis and the cytobrush’ head was cut and put into a vial with RNAlater reagent. A second cytobrush was inserted to obtain more mucosal sample, and this head was put as well into RNAlater reagent (Invitrogen). The samples were preserved at 4°C. Then, the cytobrush heads were removed and the cell pellet was obtained by centrifugation; TRIzol Reagent (Zymo) was added to lyse the cells. The samples were preserved at -80°C until RNA extraction. The anal sampling and cytology analysis was done by qualified staff from the reference lab “Laboratorio Clínico VID”.

### T cells basal activation profile

Fresh whole blood was stained for 25 min at room temperature in the dark with the monoclonal antibodies anti-CD4-PerCp-Cy5.5 clone OKT4, anti-CD8-eFluor 450 clone OKT8, anti-HLA-DR-FITC clone LN3, anti-CD69-APC clone FN50, anti-CD38-PE-Cy7 clone HIT2, and fixable viability dye-eFluor 506 (Thermo Fisher Scientific, Wilmington, DE, United States). Erythrocytes were lysed (BD FACS Lysing Solution, BD Biosciences, San Jose, CA, United States) according to the manufacturer’s instructions, next, the cells were permeabilizated and stained with anti-Ki-67-PE clone B56 (BD Biosciences, San Jose, CA, United States) and anti-CD3-Alexa eFluor 700 clone UCHT1 (Thermo Fisher Scientific, Wilmington, DE, United States) for 25 min at 4°C in the dark. The cells were fixed with 1% formaldehyde, acquired using a BD LSRFortessa™ flow cytometer and, analyzed in FlowJo software (Becton–Dickinson, San Diego, CA, USA).

### HIV-1-specific T cells responses

PBMCs were isolated by density gradient with Ficoll-Hypaque (Sigma-Aldrich, St. Louis, MO, United States), washed with Phosphate-Buffered Saline (PBS) (Lonza, Rockland, ME, United States), and suspended in RPMI medium (Lonza, Rockland, ME, United States) supplemented with fetal bovine serum (FBS) at 10% (Gibco, Grand Island, NY, United States) and penicillin/streptomycin at 1% (Thermo Fisher Scientific, Wilmington, DE, United States). The cells were culture in the presence of 3μg/mL brefeldin A, 2uM monensin, 500μg anti-CD28 clone CD28.2, 500μg anti-CD49d clone 9F10 (Thermo Fisher Scientific, Wilmington, DE, United States), and stimulated overnight with a pool of peptides from HIV-1 subtype B consensus Gag (National Institutes of Health, AIDS Research and Reference Reagents Program). *Staphylococcus* enterotoxin B (SEB) was used as the positive control. After the overnight stimulus, the supernatants were collected and stored at -80°C until the cytometric bead array (CBA) assay, and the cells were permeabilizated and stained with monoclonal antibodies CD3-Alexa eFluor 700 clone UCHT1, CD4-eFluor 660 clone OKT4, CD8-eFluor 450 clone OKT8, TNF-α-PerCp-Cy5.5 clone Mab11, fixable viability dye-eFluor 506 (Thermo Fisher Scientific, Wilmington, DE, United States), IFN-γ-brilliant Violet 711 clone 4S.B3 (BioLegend INC, San Diego, CA), Granzyme B-FITC clone GB11 and, MIP1-β-PE clone D21-1351 (BD Biosciences, San Jose, CA, United States), during 25min at 4°C in the dark. The cells were fixed with 1% formaldehyde, acquired using a BD LSRFortessa™ flow cytometer and, analyzed in FlowJo software (Becton– Dickinson, San Diego, CA, USA).

### NK-cell profile

PBMCs were stimulated with IL-12 and IL-15 (20μg/mL) for 48 hours; 3μg/mL brefeldin A and 2uM monensin were added 24 hours post culture. Cells were stained with monoclonal antibodies CD16-Alexa Fluor 647 clone 38G (BioLegend INC, San Diego, CA), CD56-PE-Cy5 clone CMSSB, NKG2D-PerCP-eFluor710 clone 1D11, and fixable viability dye-eFluor 506 (Thermo Fisher Scientific, Wilmington, DE, United States) for 25 min at room temperature in the dark. Then, the cells were permeabilized and stained with CD3-Alexa eFluor 700 clone UCHT1 (Thermo Fisher Scientific, Wilmington, DE, United States), IFN-γ -Brilliant Violet 711 clone 4SB3 (BioLegend INC, San Diego, CA), Granzyme-FITC clone GB11 and, Perforin-PE (BD Biosciences, San Jose, CA, United States), during 25min at 4°C in the dark. Finally, the cells were fixed with 1% formaldehyde and acquired using a BD LSR Fortessa™ flow cytometer and analyzed in FlowJo software (Becton–Dickinson, San Diego, CA, USA).

### mRNA quantification by real-time RT-PCR from PBMC and mucosal samples

Total RNA was purified using the Direct-zol RNA Miniprep kit (Zymo Research), treated with DNase and, retrotranscribed to cDNA using High-Capacity cDNA Reverse Transcription Kit (Thermo Fisher Scientific, Wilmington, DE, United States). PCR reactions were performed using the Maxima SYBR Green qPCR master mix kit (Fermentas). The specific primers and PCR conditions are shown in Tables 1 and 2 in S1 Text. Real-time RT-PCR was performed in a QuantStudio 5 Real-Time PCR System (Thermo Fisher Scientific, Wilmington, DE, United States). The data are expressed as mRNA relative units of each gene normalized against the constitutive gene PGK (Phosphoglycerate kinase) using the formula 1.8^−[ΔCt]^, where 1.8 corresponds to the mean PCR efficiency of 90%.

**Table 1.**
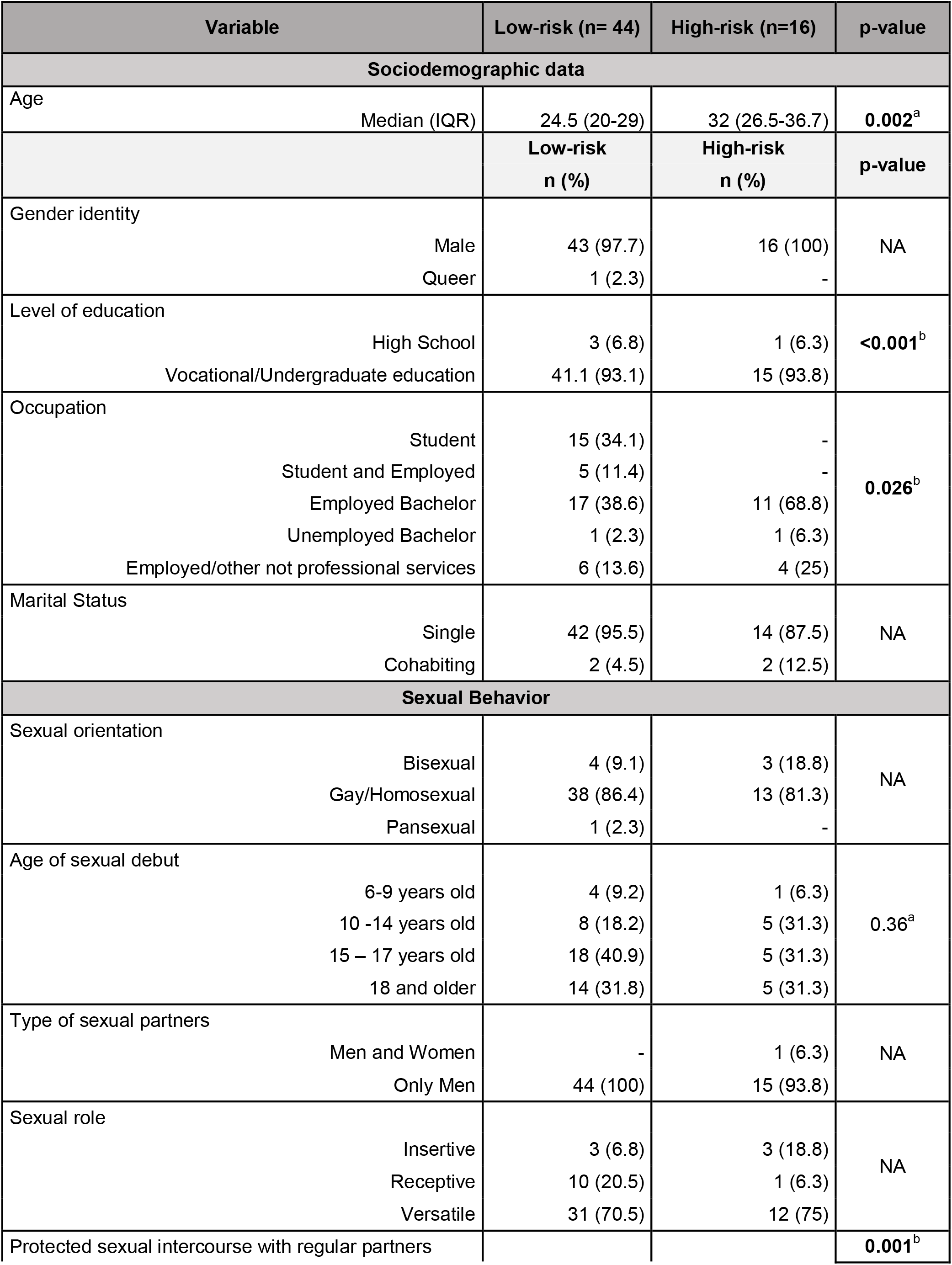
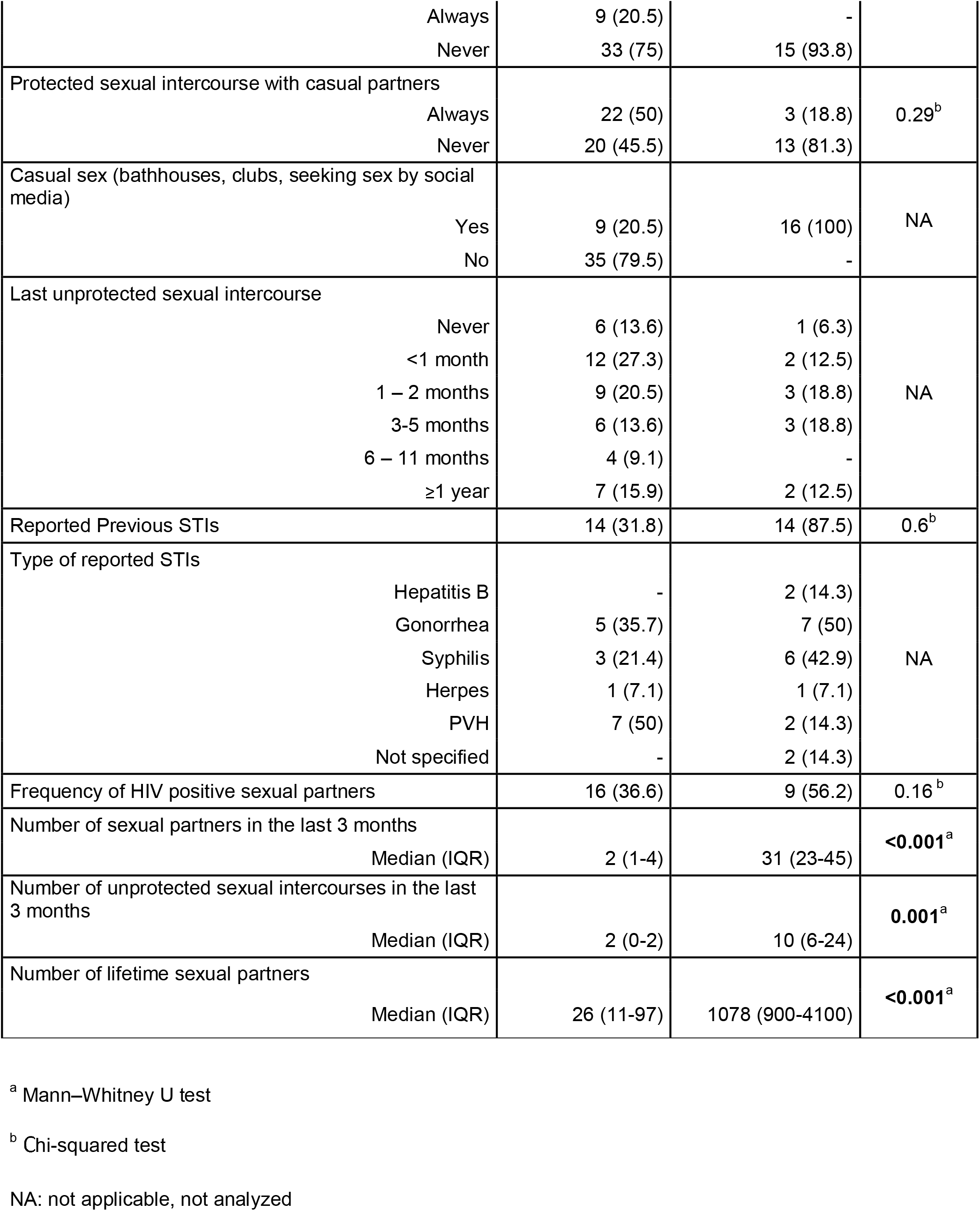
Sociodemographic and sexual behavior data of low and High-risk groups of MSM.

**Table 2.** Plasmatic levels of cytokines in both groups of MSM.

| Cytokine | Low-risk<br>(n=44) | High-risk<br>(n=16) | U Mann Whitney |
| --- | --- | --- | --- |
|  | pg/mL Median (IQR) | pg/mL Median (IQR) | p-value |
| **IL-6** | 13.9 (12.8-15.0) | 14.0 (12.9-14.9) | 0.95 |
| **IL-8** | 24.3 (23.0-26.2) | 26.1 (24.4-29.1) | 0.08 |
| **IL-10** | 15.7 (15.4-15.9) | 15.5 (15.4-15.6) | 0.23 |
| **IL-12** | 17.4 (17.0-19.0) | 17.1 (16.8-17.5) | 0.10 |
| **TNF-α** | 0.0 (0.0-4.7) | 0.0 (0.0-3.9) | 0.61 |

### Cytokines quantification by CBA

The levels of IL-1β, IL-6, IL-8, IL-12p70, TNF-α, and IL-10 were analyzed in plasma and CTL culture supernatants. The concentration of cytokines was determined by BD™ cytometric bead array (CBA) Human Inflammation Kit (Becton–Dickinson, San Diego, CA, USA), according to the manufacturer’s instructions. The data were analyzed in BD LSR Fortessa™ flow cytometer and FlowJo software (Becton–Dickinson, San Diego, CA, USA).

## Results

### The MSM from the high-risk group show risky sexual behaviors and remain HIV seronegative

The median age was 24.5 and 32 for subjects in the low and high-risk groups, respectively. In both groups, most of subjects identified themselves as gay/homosexual males, were single, and had access to vocational/professional education. In comparison with the low-risk group, the MSM in the high-risk group showed more risky sexual behaviors, including a higher number of sexual partners in the last three months (p<0.001), higher number of lifetime sexual partners (p<0.001), more unprotected intercourses in the last three months (p=0.001), and lower frequency of protected intercourses with their regular partners (p=0.001). In addition, in the high-risk group 81.3% practice receptive or versatile role; 81.3% never use condoms with their casual partners, 87.5% reported a previous history of STI, and 100% seek casual sex at bathhouses and clubs. The sociodemographic and sexual behavior data of MSM groups are described in Table 1.

### Seronegative MSM at high-risk of HIV-1 infection show a low T cell activation profile and a higher expression of Serpin A1

To explore the basal activation profile of CD4+ and CD8+ T cells, the percentage of cells expressing CD38, HLA-DR, CD69, and Ki-67 molecules was quantified by flow cytometry (Fig 1 in S1 Text). The high-risk group showed a low activation profile in T cells with lower percentage of CD4+CD38+ (p=0.002), CD4+HLADR-CD38+ (p=0.027), and CD4+Ki67+ cells (p=0.048). Likewise, this group had a higher percentage of CD4+HLADR-CD38-(p=0.013). Although it was not statistically significant, there was a tendency in the high-risk group to present a lower percentage of CD8+HLADR+CD38-cells (p=0.058). No differences were found in the expression of CD69 between both groups (Fig 2). To identify whether high-risk group exhibit differences in the basal levels of plasmatic cytokines and soluble factors, the concentrations of IL-6, IL-8 IL-10, IL-12p70, and TNF-α were quantified by CBA, and the mRNA expression of Elafin, Serpin A1, MIP1-β, RANTES, IL-1β, IL-18, IL-22, Caspase 1, and FoxP3, in PBMCs through qPCR. Compared with the low-risk group, the MSM in the high-risk group exhibit higher mRNA levels of Serpin A1 (p=0.018) and a tendency to express more MIP1-β (0.27 vs. 0.10; p=0.06). No differences were found regards the other soluble factors evaluated (Tables 2 and 3).

**Table 3.** mRNA gene expression in PBMCs of both groups of MSM.

| Gene | Low-Risk<br>(n=29) | High-Risk<br>(n=9) | U Mann-Whitney |
| --- | --- | --- | --- |
|  | mRNA relative units<br>Median (IQR) | RFU Median (IQR) | p-value |
| **Caspase 1** | 0.285 (0.0634-0.174) | 0.330 (0.174-0.534) | 0.49 |
| **Elafin** | 0.0003 (0.0001-0.0006) | 0.0005 (0.0001-0.0012) | 0.51 |
| **IL-1β** | 0.334 (0.167-0.855) | 0.334 (0.167-0.855) | 0.79 |
| **IL-18** | 0.004 (0.002-0.007) | 0.004 (0.003-0.008) | 0.82 |
| **Serpin A1** | 1.612 (0.785-2.750) | 2.326 (1.390-3.456) | **0.01** |
| **FoxP3** | 0.005 (0.001-0.012) | 0.005 (0.001-0.011) | 0.76 |
| **IL-22** | 0.0005 (0.0003-0.0012) | 0.0004 (0.0003-0.0006) | 0.74 |
| **MIP1-β** | 0.130 (0.059-0.237) | 0.245 (0.102-0.349) | 0.06 |
| **RANTES** | 1.223 (0.473-1.839) | 1.258 (0.604-2.370) | 0.15 |

The data are expressed as mRNA relative units of each gene (median and interquartile range -IQR) normalized against the constitutive gene PGK.

**Fig.1.**
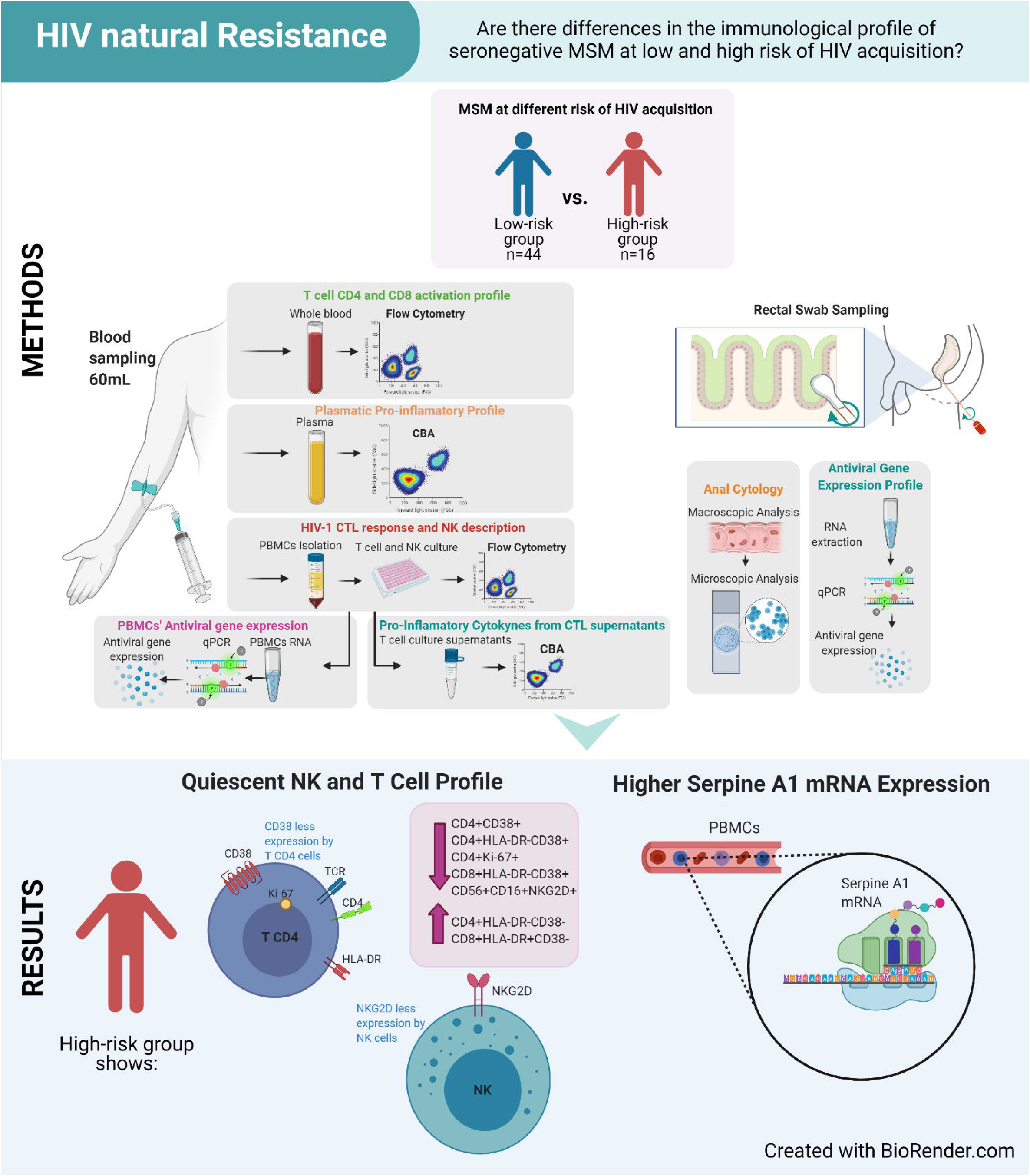
Graphic Abstract.

**Fig 2.**
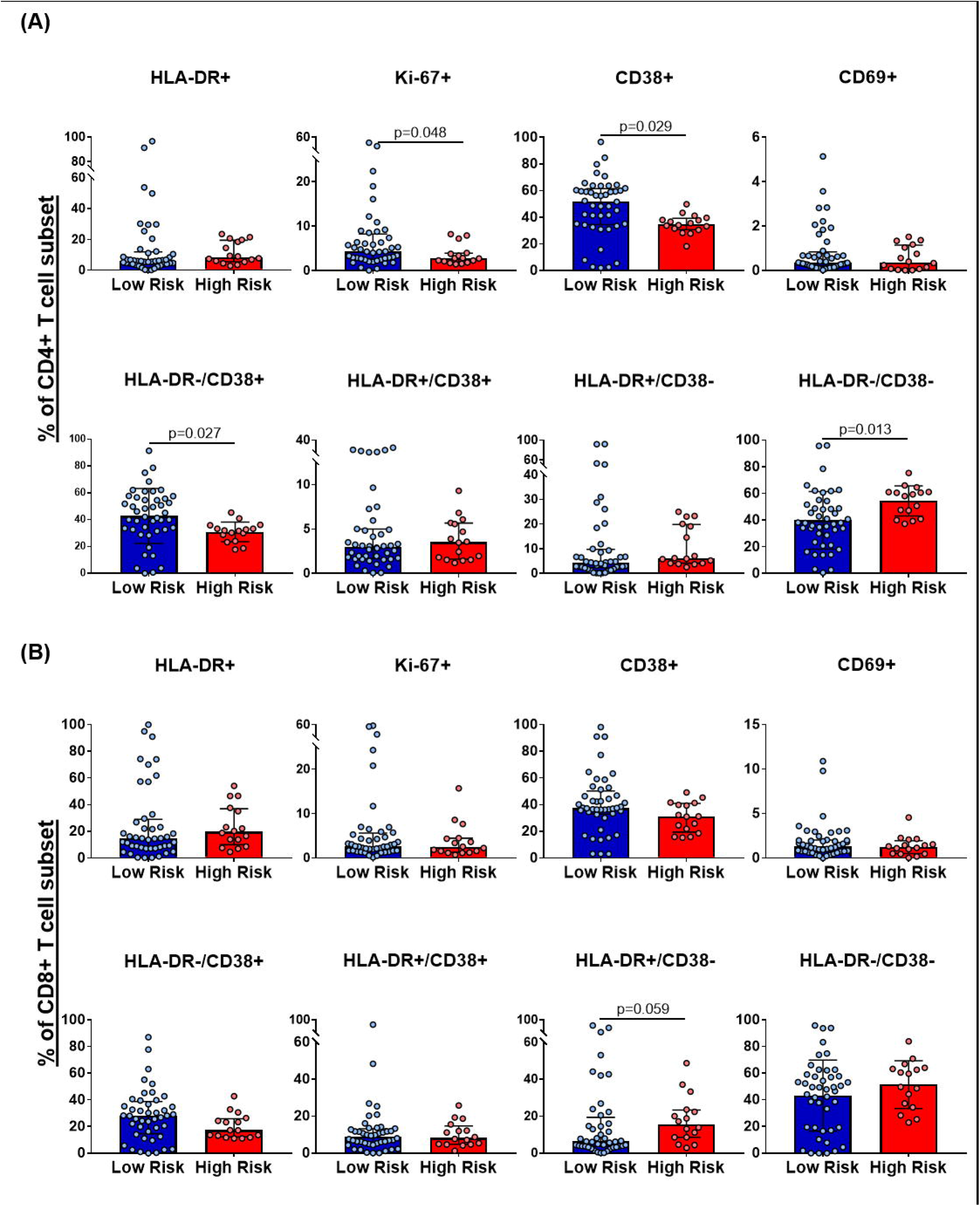
CD4+ and CD8+ T cells basal activation profile in both groups of MSM. Comparison of the percentage of **(A)** CD4+ and **(B)** CD8+ T cells expressing HLA-DR, Ki-67, CD38, and CD69 activation markers between the groups. High-risk group n=16, low-risk group n=44. All data are reported after background correction. The comparison was realized with the Mann-Whitney U test or Student T-test according to the data distribution.

### There are no differences in the T cells response to HIV-1 Gag peptides between both groups of MSM

The intracellular expression of Granzyme B, MIP1-β, TNF-α, and IFN-γ by CD4+ and CD8+ T cells was measured after stimulus with a pool of peptides from HIV-1 subtype B consensus Gag or SEB (Fig 2 in S1 Text). Likewise, the levels of IL-10, IL-12, IL-1b, IL-6, IL-8, and TNF-α in the culture’s supernatants were quantified by CBA. All the 60 MSM showed a strong specific response to SEB, however, the HIV-1 specific response only was detected in some subjects and there was not a defined pattern of response by risk group. No differences were found in the magnitude nor index of T cell polyfunctionality between the MSM groups in response to HIV-1 Gag peptides (Fig 3). Finally, no statistical differences were found between the groups in the levels of IL-10, IL-12, IL-1β IL-6, IL-8, and TNF-α in the supernatants of HIV-1 stimulated cells (Table 3 in S1 Text).

**Fig 3.**
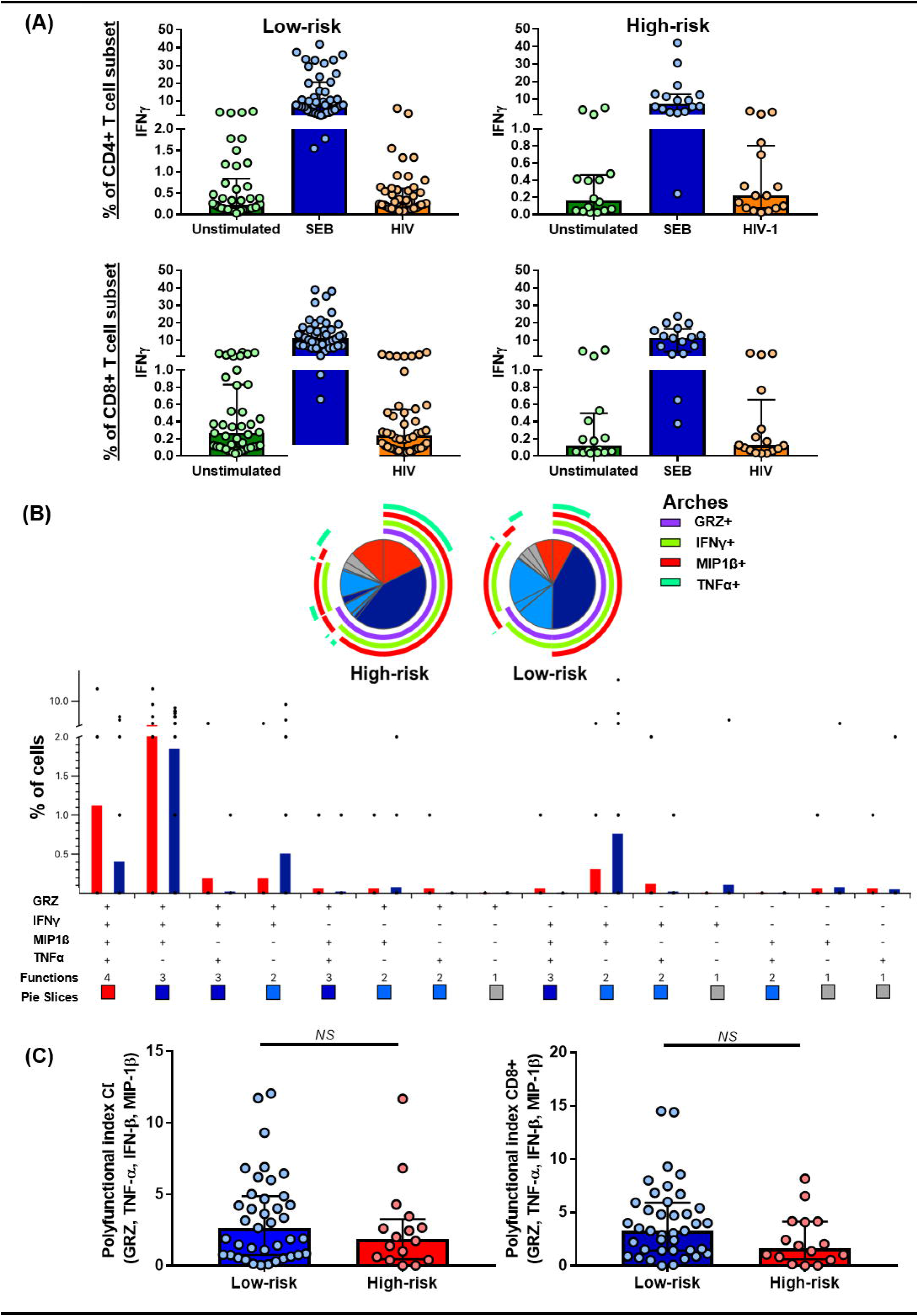
T cells response against common pathogens (SEB) and a pool of HIV-1 Gag peptides. **(A)** IFN-γ expression by unstimulated CD4+ and CD8+ T cells or after SEB and HIV-1 Gag peptides stimulus. **(B)** Comparison of the polyfunctional profiles of Gag-specific responses in CD4+ T cells from both MSM groups. The slices of the pies correspond to the proportions of Gag-specific CD4+ T cells expressing 1 (grey), 2 (light blue), 3 (dark blue), or 4 (red) functions, until n simultaneous parameters (n+1 dimensions) calculated using Boolean gating; the results are presented as mean. In the permutation analysis carried out in the SPICE platform only data higher than 0.1% were included (after background subtraction). **(C)** INDEX of polyfunctionality (pINDEX) of CD4+ and CD8+ T cells from both MSM groups based on the proportions of cells producing intracellular combinations of Granzyme B, MIP1-β, TNF-α, and IFN-γ; the pINDEX was calculated with Funky Cells software (31); the comparison was realized with the Mann-Whitney U test. For all the panels: high-risk group n=16, low-risk group n=44.

### Seronegative MSM at high-risk of HIV-1 infection show a low expression of NKG2D on NK cells

To explore the different NK cells subpopulations in both MSM groups, the CD56 and CD16 expression was analyzed in the CD3^-^ cells from fresh peripheral blood (Fig 3A in S1 Text). Both groups of MSM showed a similar distribution of NK cells subpopulations. Then, the expression of IFN-γ, Perforin, Granzyme B, and NKG2D was assessed in total NK cells from PBMCs after being stimulated for 48 hours with the combination of IL-12 and IL-15 (Fig 2 in S1 Text). The high-risk group showed a lower percentage of total NK cells expressing NKG2D (p=0.021). No differences were found in the magnitude nor polyfunctional index in those cells (Fig 4).

**Fig 4.**
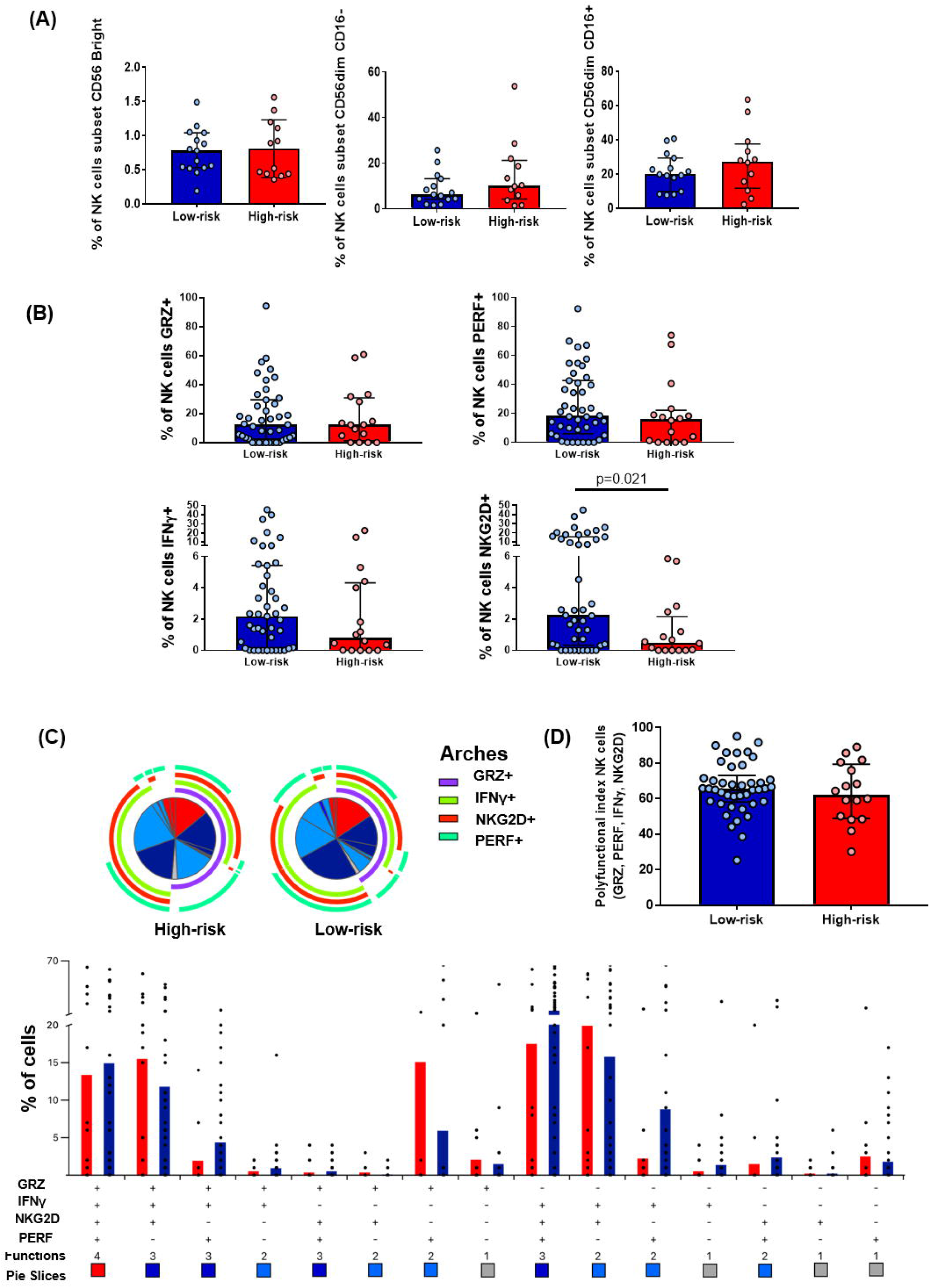
NK cells subset distribution, NKG2D expression, and polyfunctional index. (**A)** Comparison between both MSM groups of NK cells subsets (CD56bright, CD56dim CD16+, CD56dim CD16-) from fresh peripheral blood. The comparison was realized with the Mann-Whitney U test. Low-risk n=15, high-risk n=12. **(B)** Comparison of percentage of total NK cells (CD3-CD56+CD16- and CD3-CD56+CD16+ PBMCs) expressing Granzyme B, Perforin, IFN-γ or NKG2D between both groups of MSM. The comparison was realized with the Mann-Whitney U test. Data are presented after background subtraction. Low-risk n=43, high-risk n=14. **(C)** Comparison of the polyfunctional profiles of total NK cells (CD3-CD56+CD16- and CD3-CD56+CD16+ PBMCs) from both MSM groups. The slices of the pies correspond to the proportions of total NK cells expressing 1 (grey), 2 (light blue), 3 (dark blue), or 4 (red) functions, until n simultaneous parameters (n+1 dimensions) calculated using Boolean gating; the results are presented as mean. In the permutation analysis carried out in the SPICE platform only data higher than 0.1% were included (after background subtraction). **(D)** INDEX of polyfunctionality (pINDEX) of total NK cells from both MSM groups based on the proportions of cells expressing Granzyme B, Perforin, IFN-γ and NKG2D; the pINDEX was calculated with Funky Cells software; the comparison was realized with the Mann-Whitney U test. Low-risk n=43, high-risk n=14.

### There are no differences in the expression of antiviral genes in anal mucosa between both groups of MSM

To explore the macroscopic and microscopic state of anal mucosal tissue and the expression of the antiviral genes *HPN1, HBD2, HBD3, SLPI, RNAse7, TRIM5*α, and *APOBE3G*, an anal sampling was made. In both groups, most MSM presented no symptoms, a healthy anus, and a low frequency of intraepithelial lessons. No statistical differences were found regarding symptoms or the macro and microscopic state of anal mucosa between both groups (Fig 5). Quantification of the mRNA expression of the antiviral genes was no possible due to the low quantity of mRNA obtained, so a qualitative comparison of the detected genes was made between both groups. There were no statistical differences in the detection of the antiviral genes *HPN1, HBD2, HBD3, SLPI, RNAse7, TRIM5*α, and *APOBE3G* (Table 4 in S1 Text).

**Fig 5.**
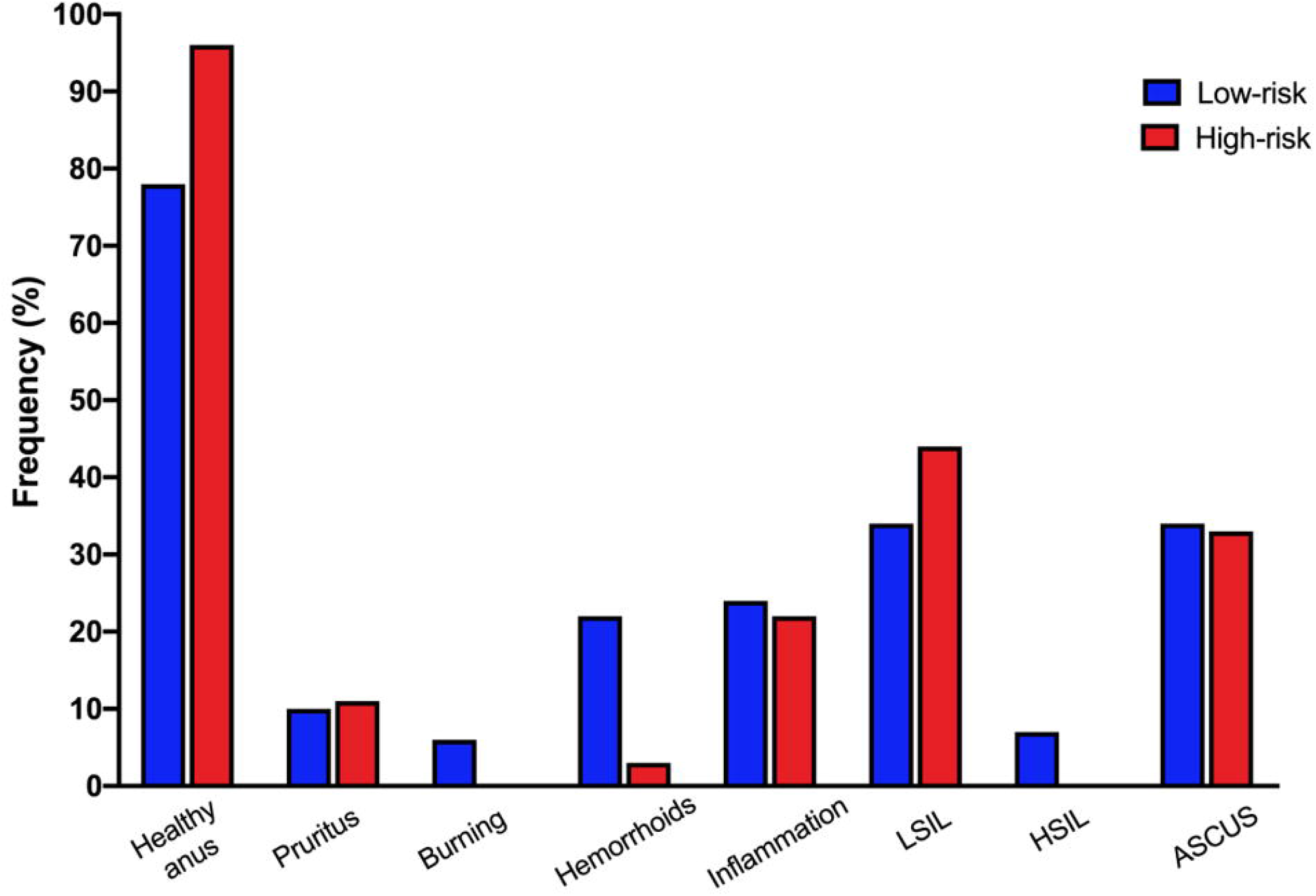
Symptoms, macroscopic and microscopic findings in anal mucosa tissue of both MSM groups. Comparison of symptoms, macroscopic, and microscopic findings in the anal mucosa tissue of both MSM groups.Data represent the percentages of people presenting each condition; there were no statistical differences between both groups through Chi-square test. Low-risk n=29, high-risk n=9. ASCUS: atypical squamous cells of uncertain significance; LISIL: low-grade squamous intraepithelial lesion; HSIL: high-grade squamous intraepithelial lesion.

## Discussion

This study reached a population with challenging access given the conditions of the vulnerability of the LGBTI population in our country (32) and the inclusion criteria that directed the recruitment of individuals with very high-risk sexual behaviors in the absence of sex work. These results and our previous study showed that MSM individuals included in the high-risk group are at extreme risk of HIV infection, showing behaviors that exceed by far the factors associated with seroconversion in the other MSM cohorts, who remain seronegative (30). Although no correlations were observed between the immunological variables and the sexual behaviors of MSM, significant differences were found between both groups regarding immune factors that have been previously associated with resistance to HIV-1 and that limit its transmission/acquisition.

The MSM group at high-risk of HIV infection showed a low activation profile of T and NK cells. The lower T cells activation profile has been described previously in HESN, which has low expression of T cell activation markers (33) and higher percentages of Treg cells (34). The study of Camara *et al*. (35) showed that HESN of serodiscordant couples had a lower percentage of CD4+ T cells expressing CD38 than control subjects. A similar low percentage of CD4+ T cells positive for CD38 has been described in MSM HESN from an Amsterdam cohort (33), in HESN from female sex workers of the Pumwani cohort in Kenya (36), as well as in Colombian elite controllers (37). Moreover, it has been described that the low percentages of T cells expressing activation molecules are related to a lower susceptibility of HIV-1 infection *in vitro*, and the persistent HIV-1 seronegative status is associated with lower T cell activation (33,38).

Our group has previously described that elite controllers exhibit a lower percentage of NK and T cells expressing activation molecules than HIV-1 progressors (39), which, together with the aforementioned studies, point to a protective quiescent cell profile characterized by a low activation of immune cells without the loss of functionality. These findings are similar to those in this cohort, where a low activation profile of T cells with a strong response to common pathogens and HIV-1 peptides was observed in some individuals. Although there were not found other studies reporting low expression of NKG2D in HESN nor elite controllers, Muntasell *et al*. demonstrated in a model of CMV infection that the exposure to the virus resulted in a decreased expression of NKG2D, which selectively limited the ability of NK cells to kill target cells expressing high levels of NKG2D ligands, while preserving the expression of other NK activation molecules and the NK cytotoxic potential (40). Likewise, we previously found that MSM at high-risk from this cohort showed higher cytotoxic capacity and IFN-γ production in response to K562 cell-stimuli compared to MSM at low-risk of HIV-1 infection (41). Those findings reinforce the theory that the quiescent immune profile is a protective factor associated with the control of HIV-1 and is not limited to T cells. To our knowledge, this is the first evidence of low expression of NKG2D by NK cells associated with natural resistance to HIV-1 infection in HESN. A higher expression of Serpin A1 in PBMCs was found in the high-risk group. High levels of Serpin A1 have previously been described in the genital mucosa of HESN female sex workers (42), genital and oral mucosa of HESN serodiscordant couples, and GALT tissue of HIV-1 elite controllers (9). The anti-HIV function of Serpin A1 in the mucosa is well understood, as it prevents damage to the tissue integrity, thus avoiding the inflammatory response and transmigration of the virus to other tissues (REF). Although less discussed, the anti-HIV effect of Serpin A1 in peripheral blood is significant considering that this serine protease: (i) inhibits neutrophil elastase, which promotes the entry of HIV-1 into the cell through its binding to gp120, and, through the cleavage that makes SDF-1 (CXCL12) and CXCR4, facilitating the binding of HIV-1 with this receptor; (ii) inhibits the formation of gp120; and (iii) inhibits the processing of p55 to p24 by protease (43). This anti-HIV effect has been previously demonstrated by the absence of HIV infection in whole blood compared to the presence of infection in lymphoid nodules under the same *in vitro* conditions (44), as well as in the easier replication of the virus in whole blood of individuals with inherited Serpin A1 deficiency, compared to the lower infection rate in whole blood of healthy controls (45). To our knowledge, this is the first evidence of increased expression of Serpin A1 by PBMCs in seronegative MSM cohorts at high risk of HIV-1 infection.

The limitations of our study rely upon small sample size, no probabilistic sampling, and the cross-sectional nature of it, which do not allow us to infer and extrapolate our findings to other populations, as well as to build an explicative and causality model of HIV-1 resistance in MSM at high-risk of HIV-1 infection.

Taken together, this suggests that Colombian MSM at high-risk could be HESN individuals and natural resistance against HIV-1 could be a combination of quiescent T and NK cells profiles and increased expression of Serpin A1 by PBMCs. It is necessary to continue the study of MSM at high-risk of exposure to HIV-1 to better understand their natural response to the virus and improve the prevention and therapeutic strategies against HIV-1 infection.

## Acknowledgments

The authors thank the volunteers who kindly participated in this study for their confidence and priceless help. We also thank the Corporación Stonewall, Consejo Consultivo LGBTI de Medellín, Alianza Social LGBTI de Medellín y Antioquia, Fundación Antioqueña de Infectología [FAI], Red de Apoyo Social de Antioquia [RASA], and the Global Fund to Fight AIDS-Enterritorio, all institution from Medellín, Colombia, for their support in the consecution of volunteers. We thank the National Institutes of Health, AIDS Research and Reference Reagents Program for the pool of peptides from HIV-1 subtype B consensus Gag. Finally, we thank Dr. Juan Camilo Sánchez-Arcila PhD, postdoctoral researcher at the University of California Merced and Dr. Martin Larsen PhD, creator of Funky Cells, for their advice in the cytometry and polyfunctional analysis of cells.

## REFERENCES

1. UNAIDS. UNAIDS Data 2019 [Internet]. Geneva; 2019. Available from: https://www.unaids.org/sites/default/files/media_asset/2019-UNAIDS-data_en.pdf

2. Burton DR. Advancing an HIV vaccine; advancing vaccinology. Nat Rev Immunol [Internet]. 2019 Feb 18;19(2):77–8. Available from: http://www.nature.com/articles/s41577-018-0103-6

3. Piacentini L, Fenizia C, Naddeo V, Clerici M. Not just sheer luck! Immune correlates of protection against HIV-1 infection. Vaccine [Internet]. 2008 Jun;26(24):3002–7. Available from: https://linkinghub.elsevier.com/retrieve/pii/S0264410X07013801

4. Horton RE, McLaren PJ, Fowke K, Kimani J, Ball TB. Cohorts for the Study of HIV 1–Exposed but Uninfected Individuals: Benefits and Limitations. J Infect Dis [Internet]. 2010 Nov;202(3):S377–81. Available from: https://academic.oup.com/jid/article-lookup/doi/10.1086/655971

5. Imagawa DT, Lee MH, Wolinsky SM, Sano K, Morales F, Kwok S, et al. Human Immunodeficiency Virus Type 1 Infection in Homosexual Men Who Remain Seronegative for Prolonged Periods. N Engl J Med [Internet]. 1989 Jun;320(22):1458–62. Available from: http://www.nejm.org/doi/abs/10.1056/NEJM198906013202205

6. Ranki A, Mattinen S, Yarchoan R, Broder S, Ghrayeb J, Lähdevirta J, et al. T-cell response towards HIV in infected individuals with and without zidovudine therapy, and in HIV-exposed sexual partners. Aids [Internet]. 1989 Feb;3(2):63–70. Available from: http://journals.lww.com/00002030-198902000-00002

7. Gupta A, Padh H. The global distribution of CCR5 delta 32 polymorphism: role in HIV-1 protection. BMC Infect Dis [Internet]. 2012 Dec 4;12(S1):O16. Available from: https://bmcinfectdis.biomedcentral.com/articles/10.1186/1471-2334-12-S1-O16

8. Rodríguez-Mora S, De Wit F, García-Perez J, Bermejo M, López-Huertas MR, Mateos E, et al. The mutation of Transportin 3 gene that causes limb girdle muscular dystrophy 1F induces protection against HIV-1 infection. Emerman M, editor. PLOS Pathog [Internet]. 2019 Aug 29;15(8):e1007958. Available from: https://dx.plos.org/10.1371/journal.ppat.1007958

9. Gonzalez SM, Taborda NA, Feria MG, Arcia D, Aguilar-Jiménez W, Zapata W, et al. High Expression of Antiviral Proteins in Mucosa from Individuals Exhibiting Resistance to Human Immunodeficiency Virus. Landay A, editor. PLoS One [Internet]. 2015 Jun 19;10(6):e0131139. Available from: https://dx.plos.org/10.1371/journal.pone.0131139

10. Scott-Algara D, Truong LX, Versmisse P, David A, Luong TT, Nguyen N V, et al. Cutting edge: increased NK cell activity in HIV-1-exposed but uninfected Vietnamese intravascular drug users. J Immunol. 2003 Dec;171(11):5663–7.

11. Habegger de Sorrentino A, Sinchi JL, Marinic K, López R, Iliovich E. KIR-HLA-A and B alleles of the Bw4 epitope against HIV infection in discordant heterosexual couples in Chaco Argentina. Immunology [Internet]. 2013 Oct;140(2):273–9. Available from: http://doi.wiley.com/10.1111/imm.12137

12. Pallikkuth S, Wanchu A, Bhatnagar A, Sachdeva RK, Sharma M. Human immunodeficiency virus (HIV) gag antigen-specific T-helper and granule-dependent CD8 T-cell activities in exposed but uninfected heterosexual partners of HIV type 1-infected individuals in North India. Clin Vaccine Immunol. 2007 Sep;14(9):1196–202.

13. Flórez-Álvarez L, Hernandez JC, Zapata W. NK Cells in HIV-1 Infection: From Basic Science to Vaccine Strategies. Front Immunol [Internet]. 2018 Oct 17;9. Available from: https://www.frontiersin.org/article/10.3389/fimmu.2018.02290/full

14. Mwatelah R, Mckinnon LR, Baxter C, Abdool Karim Q, Abdool Karim SS. Mechanisms of sexually transmitted infection induced inflammation in women: implications for <scp>HIV</scp> risk. J Int AIDS Soc [Internet]. 2019 Aug 30;22(S6). Available from: https://onlinelibrary.wiley.com/doi/abs/10.1002/jia2.25346

15. Kaul R, Prodger J, Joag V, Shannon B, Yegorov S, Galiwango R, et al. Inflammation and HIV Transmission in Sub-Saharan Africa. Curr HIV/AIDS Rep [Internet]. 2015 Jun 16;12(2):216–22. Available from: http://link.springer.com/10.1007/s11904-015-0269-5

16. Appay V, Kelleher AD. Immune activation and immune aging in HIV infection. Curr Opin HIV AIDS [Internet]. 2016 Mar;11(2):242–9. Available from: http://journals.lww.com/01222929-201603000-00016

17. Appay V, Sauce D. Immune activation and inflammation in HIV-1 infection: causes and consequences. J Pathol [Internet]. 2008 Jan;214(2):231–41. Available from: http://doi.wiley.com/10.1002/path.2276

18. Fulcher JA, Romas L, Hoffman JC, Elliott J, Saunders T, Burgener AD, et al. Highly Human Immunodeficiency Virus-Exposed Seronegative Men Have Lower Mucosal Innate Immune Reactivity. AIDS Res Hum Retroviruses. 2017 Aug;33(8):788–95.

19. Card CM, Ball TB, Fowke KR. Immune quiescence: a model of protection against HIV infection. Retrovirology. 2013 Nov;10:141.

20. Yao X-D, Omange RW, Henrick BM, Lester RT, Kimani J, Ball TB, et al. Acting locally: innate mucosal immunity in resistance to HIV-1 infection in Kenyan commercial sex workers. Mucosal Immunol [Internet]. 2014 Mar 26;7(2):268–79. Available from: http://www.nature.com/articles/mi201344

21. Taborda NA, Hernandez JC, Lajoie J, Juno JA, Kimani J, Rugeles MT, et al. Short Communication: Low Expression of Activation and Inhibitory Molecules on NK Cells and CD4(+) T Cells Is Associated with Viral Control. AIDS Res Hum Retroviruses. 2015 Jun;31(6):636–40.

22. Hernandez JC, St Laurent G, Urcuqui-Inchima S. HIV-1-exposed seronegative individuals show alteration in TLR expression and pro-inflammatory cytokine production ex vivo: An innate immune quiescence status? Immunol Res [Internet]. 2016 Feb;64(1):280–90. Available from: http://link.springer.com/10.1007/s12026-015-8748-8

23. Card CM, McLaren PJ, Wachihi C, Kimani J, Plummer FA, Fowke KR. Decreased Immune Activation in Resistance to HIV 1 Infection Is Associated with an Elevated Frequency of CD4 + CD25 + FOXP3 + Regulatory T Cells. J Infect Dis. 2009 May;199(9):1318–22.

24. Abdulhaqq SA, Zorrilla C, Kang G, Yin X, Tamayo V, Seaton KE, et al. HIV-1-negative female sex workers sustain high cervical IFNLJ, low immune activation, and low expression of HIV-1-required host genes. Mucosal Immunol [Internet]. 2016 Jul 11;9(4):1027–38. Available from: http://www.nature.com/articles/mi2015116

25. McLaren PJ, Ball TB, Wachihi C, Jaoko W, Kelvin DJ, Danesh A, et al. HIV Exposed Seronegative Commercial Sex Workers Show a Quiescent Phenotype in the CD4 + T Cell Compartment and Reduced Expression of HIV Dependent Host Factors. J Infect Dis. 2010 Nov;202(3):S339–44.

26. Biasin M, Caputo S Lo, Speciale L, Colombo F, Racioppi L, Zagliani A, et al. Mucosal and Systemic Immune Activation Is Present in Human Immunodeficiency Virus–Exposed Seronegative Women. J Infect Dis [Internet]. 2000 Nov;182(5):1365–74. Available from: https://academic.oup.com/jid/article-lookup/doi/10.1086/315873

27. Saulle I, Biasin M, Gnudi F, Rainone V, Ibba SV, Caputo S Lo, et al. Short Communication: Immune Activation Is Present in HIV-1-Exposed Seronegative Individuals and Is Independent of Microbial Translocation. AIDS Res Hum Retroviruses [Internet]. 2016 Feb;32(2):129–33. Available from: http://www.liebertpub.com/doi/10.1089/aid.2015.0019

28. Tomescu C, Seaton KE, Smith P, Taylor M, Tomaras GD, Metzger DS, et al. Innate activation of MDC and NK cells in high-risk HIV-1-exposed seronegative IV-drug users who share needles when compared with low-risk nonsharing IV-drug user controls. J Acquir Immune Defic Syndr. 2015 Mar;68(3):264–73.

29. Templeton DJ, Jin F, Mao L, Prestage GP, Donovan B, Imrie J, et al. Circumcision and risk of HIV infection in Australian homosexual men. AIDS [Internet]. 2009 Nov;23(17):2347–51. Available from: http://journals.lww.com/00002030-200911130-00012

30. Ossa-Giraldo AC, Correa JS, Moreno CL, Blanquiceth Y, Flórez-Álvarez L, Contreras-Ramírez K, et al. Sexual behaviors and factors associated with condomless sexual practice in Colombian MSM at high risk of HIV transmission. Arch Sex Behav.

31. Boyd A, Almeida JR, Darrah PA, Sauce D, Seder RA, Appay V, et al. Correction: Pathogen-Specific T Cell Polyfunctionality Is a Correlate of T Cell Efficacy and Immune Protection. PLoS One [Internet]. 2015 Sep 14;10(9):e0138395. Available from: https://dx.plos.org/10.1371/journal.pone.0138395

32. Lopez Solano H. EL MOVIMIENTO LGBT EN COLOMBIA: LA CONSTRUCCIÓN DEL DERECHO DESDE ABAJO [Internet]. Universidad Santo Tomás; 2017. Available from: https://repository.usta.edu.co/bitstream/handle/11634/3942/Lopezhernan2017.pdf?s equence=1&isAllowed=y

33. Koning FA, Otto SA, Hazenberg MD, Dekker L, Prins M, Miedema F, et al. Low-Level CD4 + T Cell Activation Is Associated with Low Susceptibility to HIV-1 Infection. J Immunol [Internet]. 2005 Nov 1;175(9):6117–22. Available from: http://www.jimmunol.org/lookup/doi/10.4049/jimmunol.175.9.6117

34. Pattacini L, Baeten JM, Thomas KK, Fluharty TR, Murnane PM, Donnell D, et al. Regulatory T-Cell Activity But Not Conventional HIV-Specific T-Cell Responses Are Associated With Protection From HIV-1 Infection. JAIDS J Acquir Immune Defic Syndr [Internet]. 2016 Jun;72(2):119–28. Available from: http://journals.lww.com/00126334-201606010-00001

35. Camara M, Dieye TN, Seydi M, Diallo AA, Fall M, Diaw PA, et al. Low Level CD4 + T Cell Activation in HIV Exposed Seronegative Subjects: Influence of Gender and Condom Use. J Infect Dis [Internet]. 2010 Mar 15;201(6):835–42. Available from: https://academic.oup.com/jid/article-lookup/doi/10.1086/651000

36. Card CM, McLaren PJ, Wachihi C, Kimani J, Plummer FA, Fowke KR. Decreased Immune Activation in Resistance to HIV 1 Infection Is Associated with an Elevated Frequency of CD4 + CD25 + FOXP3 + Regulatory T Cells. J Infect Dis [Internet]. 2009 May;199(9):1318–22. Available from: https://academic.oup.com/jid/article-lookup/doi/10.1086/597801

37. Gonzalez SM, Taborda NA, Correa LA, Castro GA, Hernandez JC, Montoya CJ, et al. Particular activation phenotype of T cells expressing HLA-DR but not CD38 in GALT from HIV-controllers is associated with immune regulation and delayed progression to AIDS. Immunol Res [Internet]. 2016 Jun 2;64(3):765–74. Available from: http://link.springer.com/10.1007/s12026-015-8775-5

38. Kuebler PJ, Mehrotra ML, Shaw BI, Leadabrand KS, Milush JM, York VA, et al. Persistent HIV Type 1 Seronegative Status Is Associated With Lower CD8 + T-Cell Activation. J Infect Dis [Internet]. 2016 Feb 15;213(4):569–73. Available from: https://academic.oup.com/jid/article-lookup/doi/10.1093/infdis/jiv425

39. Taborda NA, Hernández JC, Lajoie J, Juno JA, Kimani J, Rugeles MT, et al. Short Communication: Low Expression of Activation and Inhibitory Molecules on NK Cells and CD4 + T Cells Is Associated with Viral Control. AIDS Res Hum Retroviruses [Internet]. 2015 Jun;31(6):636–40. Available from: http://www.liebertpub.com/doi/10.1089/aid.2014.0325

40. Muntasell A, Magri G, Pende D, Angulo A, López-Botet M. Inhibition of NKG2D expression in NK cells by cytokines secreted in response to human cytomegalovirus infection. Blood [Internet]. 2010 Jun 24;115(25):5170–9. Available from: https://ashpublications.org/blood/article/115/25/5170/27093/Inhibition-of-NKG2D-expression-in-NK-cells-by

41. Flórez-Álvarez L, Blanquiceth Y, Ramírez K, Ossa-Giraldo AC, Velilla PA, Hernandez JC, et al. NK Cell Activity and CD57+/NKG2Chigh Phenotype Are Increased in Men Who Have Sex With Men at High Risk for HIV. Front Immunol [Internet]. 2020 Sep 11;11. Available from: https://www.frontiersin.org/article/10.3389/fimmu.2020.537044/full

42. Burgener A, Rahman S, Ahmad R, Lajoie J, Ramdahin S, Mesa C, et al. Comprehensive Proteomic Study Identifies Serpin and Cystatin Antiproteases as Novel Correlates of HIV-1 Resistance in the Cervicovaginal Mucosa of Female Sex Workers. J Proteome Res [Internet]. 2011 Nov 4;10(11):5139–49. Available from: https://pubs.acs.org/doi/abs/10.1021/pr200596r

43. Congote LF. Serpin A1 and CD91 as host instruments against HIV-1 infection: Are extracellular antiviral peptides acting as intracellular messengers? Virus Res [Internet]. 2007 May;125(2):119–34. Available from: https://linkinghub.elsevier.com/retrieve/pii/S0168170206003972

44. Pantaleo G, Graziosi C, Demarest JF, Butini L, Montroni M, Fox CH, et al. HIV infection is active and progressive in lymphoid tissue during the clinically latent stage of disease. Nature [Internet]. 1993 Mar;362(6418):355–8. Available from: http://www.nature.com/articles/362355a0

45. Shapiro L, H.B. Pott G, H. Ralston A. Alpha 1 antitrypsin inhibits human immunodeficiency virus type 1. FASEB J [Internet]. 2001 Jan;15(1):115–22. Available from: https://onlinelibrary.wiley.com/doi/abs/10.1096/fj.00-0311com

